# Fragile X messenger ribonucleoprotein modulates IL-6 induction and inflammatory cell death in macrophages

**DOI:** 10.64898/2026.03.30.712626

**Authors:** Brittany N Macha, Chi G Weindel, Tracy Fischer, Robert O Watson, Lindsey A Ho, Krystal J Vail

## Abstract

RNA-binding proteins are key players in determining the fate of mRNA. One such RNA binding protein, Fragile X messenger ribonucleoprotein (FMRP), has an established role in RNA transcription, metabolism, translation, and degradation in the brain and reproductive system. Although FMRP is expressed in immune cells, little is known about how FMRP influences immune cell mRNA transcript outcomes. Here, we show that macrophage infection with the intracellular pathogen *Listeria monocytogenes* induces FMRP translocation from the cytoplasm to the nucleus. We show that infected macrophages lacking FMRP have impaired *Il6* induction in response to *L. monocytogenes* infection. Finally, we show that macrophages lacking FMRP have increased susceptibility to inflammatory cell death. Together, these data implicate FMRP in modulating proinflammatory gene expression during bacterial infection.

## INTRODUCTION

Cytokine production and direct killing by innate immune cells are among the first responses in the host defense against invading pathogens. This response is carefully regulated to balance effective pathogen control while mitigating the cytopathic effects of the response to host tissues (1). Immune cells respond to and engage with rapidly changing environmental pressures by altering their transcriptome and proteome. Genetic information stored in DNA requires a regimented process to generate a final protein product, including transcription to mRNA. Eukaryotic mRNAs transition through several stages including transcription, processing, nuclear export, translation to protein, and ultimately decay. Due to their potent nature, cytokines are controlled at multiple points in their lifecycle, from gene expression, post transcriptional and post translational modifications, and protein degradation. RNA binding proteins (RBP) play a critical role in determining the fate of mRNA, and specific sets of RBPs bind to mRNAs during each stage of the RNA lifecycle to form messenger ribonucleoprotein complexes. Through these complexes, RBPs govern gene expression in response to environmental stimuli by altering quantitative and qualitative aspects of mRNA transcripts, including mRNA sequence, activity, stability, location, and translation efficiency (2).

Fragile X messenger ribonucleoprotein (FMRP) is an RBP that functions in RNA metabolism, transcription, and as a translational repressor (3–5). Loss of FMRP due to genetic silencing results in Fragile X syndrome, a form of intellectual disability. Under normal circumstances, FMRP acts as a translational brake by halting ribosomes on target mRNAs, thus pausing protein translation and controlling protein concentrations (6). This is particularly important in the developing brain, which is dependent upon controlled synthesis of new proteins for development.

While the highest expression of FMRP occurs in the brain, it is expressed ubiquitously throughout the body. Consequently, impaired FMRP has clinical implications in extracerebral sites. As an example, the *FMR1* gene, which encodes FMRP, contains up to 50 CGG trinucleotide repeats under normal circumstances. Premutation occurs when trinucleotide repeats exceed normal levels, generally up to 200 CGG repeats, which results in toxic accumulation of *FMR1* mRNA rather than its genetic silencing. Clinically, premutation is associated with conditions such as premature ovarian failure in females and Fragile X-associated tremor/ataxia syndrome, most commonly manifested in males (7). The diversity of conditions associated with *FMR1* mutation and FMRP dysfunction warrant further understanding of its role in other tissues and cell types.

Although most investigations of FMRP have centered on its function in neuron plasticity and neurodevelopment, mounting evidence implicates FMRP in innate immunity, including both the actions of the native protein as well as immune dysfunction in its absence. Intriguingly, disparate phenotypes are reported amongst those with full mutations resulting in *FMR1* silencing compared with premutation carriers in which *FMR1* mRNA increases to toxic levels leading to mitochondrial dysfunction (7, 8). For example, female premutation carriers are reported to have higher incidence of autoinflammatory and autoimmune disorders and decreased cytokine production compared with age matched controls (9). In contrast, there is a decreased risk of autoimmune disorders in full mutation carriers, who in the absence of FMRP instead demonstrate an increased risk of infectious disease such as viral enteritis, pneumonia, and candidiasis (10). Indeed, FMRP has been shown to influence bacterial susceptibility (11) as well as viral replication and infectivity by interacting with viral nucleic acids and products (12–15). Furthermore, FMRP modulates oxidative stress (16), cytokine production (9, 17, 18), programed cell death (19–21), and exosome cargo loading in response to inflammation (22). Although these studies clearly implicate FMRP in susceptibility to infection, how it contributes to regulating the immune response remains unknown.

FMRP’s function in regulating translation and modulating protein levels has been well characterized (6, 23). Recent literature indicates that FMRP is also involved in transcription in addition to RNA metabolism in epithelial and mesenchymal cell types (24). Thus, we hypothesized that FMRP influences transcriptional responses to bacterial infection.

In this study, we sought to examine the function of FMRP on antibacterial immune responses using *Listeria monocytogenes* as a model intracellular pathogen. We hypothesized that FMRP levels would be dynamic in response to macrophage infection and that the absence of FMRP would alter innate immune responses following infection. We found that in infected macrophages, FMRP translocated from the cytoplasm to the nucleus and *Fmr1* transcript levels transiently increased. We observed that in the absence of FMRP, *L. monocytogenes-*induced IL-6 production was impaired, and that inflammatory cell death was enhanced in response to NLRP3 activation. These findings implicate FMRP in modulating innate immune gene expression in response to bacterial infection.

## METHODS

### Animals

Tulane National Biomedical Research Center and Tulane University are accredited by AAALAC International. All animal experiments for this study were reviewed and approved by the Tulane University Institutional Animal Care and Use Committee and all animals were cared for in accordance with the Guide for the Care and Use of Laboratory Animals. Mice were group housed by sex on ventilated racks in temperature controlled rooms on a 12 hour light/dark cycle and provided *ad libitum* food and water. Male *Fmr1*^-/y^ mice (B6.129P2-*Fmr1*^*tm1Cgr*^/J, RRID:IMSR_JAX:003025) were purchased from The Jackson Laboratory (Bar Harbor, ME) and bred to *Fmr1*^+/+^ female mice to generate *Fmr1*^+/-^ female mice. Breeding pairs consisted of female *Fmr1*^+/-^ and male *Fmr1*^+/y^ mice, and male littermates were used for experimental groups.

### Cell Culture

Bone marrow derived macrophages were differentiated from bone marrow cells isolated by washing mouse femurs with 10 ml DMEM. Harvested cells were centrifuged for 5 min at 125xg and resuspended in complete media (DMEM, 20% FBS, 1 mM sodium pyruvate, 10% MCSF conditioned media). Cells were counted, seeded at 5×10^6^ in 15 cm non-TC treated plates in 30 ml complete media, and on day 3 an additional 15 ml of media was added. Upon visual confirmation of differentiation (approximately day 7), cells were harvested with 0.04% EDTA in PBS.

### Bacterial strains and infections

Low passage glycerol lab stocks of *L. monocytogenes* strain 10304s were streaked onto brain-heart infusion (BHI) agar plates and incubated at 37°C overnight. Approximately 5 colony forming units (CFU) of *L. monocytogenes* were inoculted into 5 ml of BHI broth and cultured under static conditions overnight at 30°C, then 500 µl of the culture was subcultured into 5 ml of fresh BHI for approximately 3 hours shaken at 37°C until reaching log phase of replication. One ml of subcultured *L. monocytogenes* was centrifuged at 1950xg for 3 min at 25°C. The supernatant was discarded, the pellet washed twice with 1 ml PBS and the concentration of bacteria was determined spectrophotometrically at an optical density of 600 nm (OD600) where OD 1.0 represents approximately 1×10^9^. Bacterial concentrations were verified by plating the inoculum and counting CFUs. Bacterial suspensions were diluted to the desired concentration. BMDMs were seeded in 12-well dishes at 5×10^5^ cells/well and allowed to adhere overnight. Macrophages were infected with *L. monocytogenes* at a MOI of 5 for gene expression and immunofluorescent staining studies, MOI of 1 for CFU experiments, or MOI of 10 for immunoblot analysis, or mock-infected with a matching volume of media. Following infection, macrophages were centrifuged for 10 minutes at 1000 rpm then incubated at 37°C for 30 minutes. After 30 minutes of incubation, the media was removed and monolayers were washed with gentamycin to kill non-internalized bacteria. Complete media was replaced and cells were cultured at 37°C until the indicated time point. Cell supernatants were collected and immediately frozen for cytokine analysis.

### Cell stimulations

Primary BMDMs were seeded in 12-, 24-, or 96-well plates at a density of 5×10^5^, 2.5×10^5^ or 5×10^4^ cells per well, respectively, and allowed to adhere overnight. Interferon stimulated genes (ISG) and proinflammatory genes were induced by stimulating cells with 100 ng/ml of LPS or transfecting 1µg/ml of poly I:C (InvivoGen) using Lipofectamine 2000 (Thermo Fisher) for 4 hours. For signal 1 cell death experiments, cells were primed with 100 ng/ml of LPS or 10 ng/ml of Pam3CSK4. For activation of the NLRP3 inflammasome, cells were primed with 20 ng/ml LPS or 10 ng/ml of Pam3CSK4 for 3h, followed by the stimulating with 10 µM nigericin for the indicated timepoints. AIM2 inflammasome was activated by priming cells with 20 ng/ml LPS or 10 ng/ml Pam3CSK4 for 3h followed by transfecting 1 µg/ml poly(dA:dT) using Lipofectamine 200 for 4 hours. NOD2/NLRP3 was activated by priming cells with 20 ng/ml LPS or 10 ng/ml Pam3CSK4 for 3h followed by transfecting 25 ng/ml MDP for the indicated time points. At the indicated time points, cell supernatants were collected and immediately used for LDH analysis. Cell lysates were harvested and frozen in radioimmunoprecipitation assay (RIPA) lysis buffer for immunoblot analysis or Trizol for RTqPCR. Cell images for PI cell death analysis were obtained on a Zeiss Axio Observer fluorescent microscope.

### LDH Cytotoxicity Assay

Macrophage supernatant samples were harvested at 0, 2, and 4 hours post stimulation. LDH was measured using the CyQUANT LDH Cytotoxicity Assay Kit (Invitrogen) according to the manufacturer’s instructions. Cytotoxicity was calculated using the following formula: % cytotoxicity = 100 x ((experimental LDH release – spontaneous LDH release) / (maximum LDH release – spontaneous LDH release)).

### Quantitative RT-PCR

Trizol reagent (Invitrogen) was used for total RNA extraction according to the manufacturer’s protocol. RNA was isolated using Direct-zol RNAeasy kits (Zymo Research). cDNA was synthesized with Maxima First Strand cDNA synthesis kit (Thermofisher) according to the manufacturer’s protocol. RT-qPCR was performed in triplicate wells using PowerUp SYBR Green Master Mix. Data were analyzed on a QuantStudio 6 Real-Time PCR System (Applied Biosystems).

### Immunoblot

Cell monolayers were washed with 1X PBS and lysed in 1X RIPA lysis buffer (150 mM NaCl, 1.0% NP-40, 0.5% sodium deoxycholate, 0.1% SDS, 50 mM Tris, pH 8.0) with protease and phosphatase inhibitors (1 tablet per 10 ml; Pierce). DNA was degraded using 250 units benzonase (Milipore Sigma E1014). Samples were boiled at 95°C for 5 minutes. Proteins were separated by SDS-PAGE and transferred to PDVF membranes. Membranes were blocked for 1 hour at room temperature in 5% Bovine Serum Albumin (BSA) and incubated overnight at room temperature with the following antibodies: Tubulin 1:1000 (Abcam 179513, RRID:AB_3073861), FMRP 1:1000 (Abcam ab17722; RRID:AB_2278530). Membranes were washed 3x in 1X TBS 0.1% Tween 20 and incubated with appropriate secondary antibody (Abcam) for 1 hour at RT (20°C) prior to imaging on an iBright Imaging System (Invitrogen).

### Immunofluorescence microscopy

For fixed cells, BMDMs were seeded at 2×10^5^ cells/well on glass coverslips in 24-well dishes. At the indicated time points, cells were fixed in 4% paraformaldehyde for 10 min at 37°C, then washed three times with PBS. Coverslips were incubated at room temperature in primary antibody (FMRP, Abcam ab17722; RRID:AB_2278530) diluted in TBS+0.25% Triton-X + 5% Normal Goat Serum for 3h. Cell monolayers were then washed three times in PBS and incubated in secondary antibody for 1h. Coverslips were incubated with DAPI for 5 minutes, then washed twice with PBS and mounted on glass slides for imaging. Images were obstained using a Leica SP8 Confocal microscope equipped with a 60X oil immersion objective. Maximum intensity projections of z-stacks were obtained and projected images were thresholded such that FMRP in macrophages was masked and quantified. For live cell imaging, cells were seeded in 96-well black clear bottom plates (Cellvis, #P96-1.5H-N) and treated as described in Cell Stimulations. Cells were treated with 1 µg/ml propidium iodide and 2 drops/ml of NucBlue Live Ready Probe Reagent (Invitrogen). Images were obtained using a Zeiss Axio Observer inverted fluorescent microscope and analyzed using ImageJ.

### IL-6 ELISA

Macrophage supernatant samples were harvested at 0-, 3-, and 6 hours post *L. monocytogenes* infection and stored at −80°C until thawing on the day of the assay. A mouse IL-6 ELISA kit (Invitrogen, cat# KMC0061) was used according to the manufacturer’s instructions and absorbance read on a GloMax Explorer (Promega) plate reader.

### Statistical analysis

All data are representative of at least 2 independent experiments with n ≥ 3. Except where indicated in the figure legend, due to small sample sizes, exact *p*-values were conducted to assess statistical significance between groups, which did not rely on large-sample approximations inherent in parametric methods such as the classic t-test. We assumed independence among BMDM samples and exchangeability between groups. We accepted a larger Type I error resulting from the smallest sample scenario of 20 possibilities derived from 3 BMDM samples per group, thereby determining significance at an alpha level of 0.10.

In order to quantify the association between FMRP expression area and cellular region in mouse BMDMs, we constructed a complete factorial experiment within a linear regression model comprising FMRP expression area and cellular region (cytosolic and nuclear) as the dependent and independent variables, respectively. We also included *L. monocytogenes* infection status in the model and a cellular region by infection status interaction term. Thus, we statistically accounted for these additional variables, which may affect the primary relationship of interest. We assessed the overall and cellular region specific *p*-values of the infection effect using appropriate contrasts comprising linear combinations of the regression parameters.

Exact and linear regression *p*-values were computed in Python version 3.13.9 and SAS version 9.4, respectively.

## RESULTS

### *Listeria monocytogenes* induces FMRP translocation to the nucleus

RNA binding proteins (RBP) such as FMRP modulate gene expression in various ways, including regulation of transcription, processing, nuclear export, and translation. Endogenous FMRP is predominantly localized to the cytoplasm (25) in multiple subcellular compartments including perinuclear regions, and mitochondria, as well as to a lesser extent in the nucleolus (26). Consistent with these reports, we observed a wide cytoplasmic distribution of FMRP in resting murine bone marrow derived macrophages (BMDMs) by fluorescence microscopy (Fig 1A upper panels). We next wished to identify how infection impacts FMRP subcellular localization. To assess the impact of infection on the subcellular localization of FMRP, we infected mouse BMDMs with green fluorescent protein (GFP)-tagged *L. monocytogenes*, a gram-positive, facultative intracellular bacterium. Using fluorescence microscopy, we observed that FMRP was enriched in the nucleus in infected cells within 4 hours post-infection (Fig 1A lower panels).

**Figure 1:**
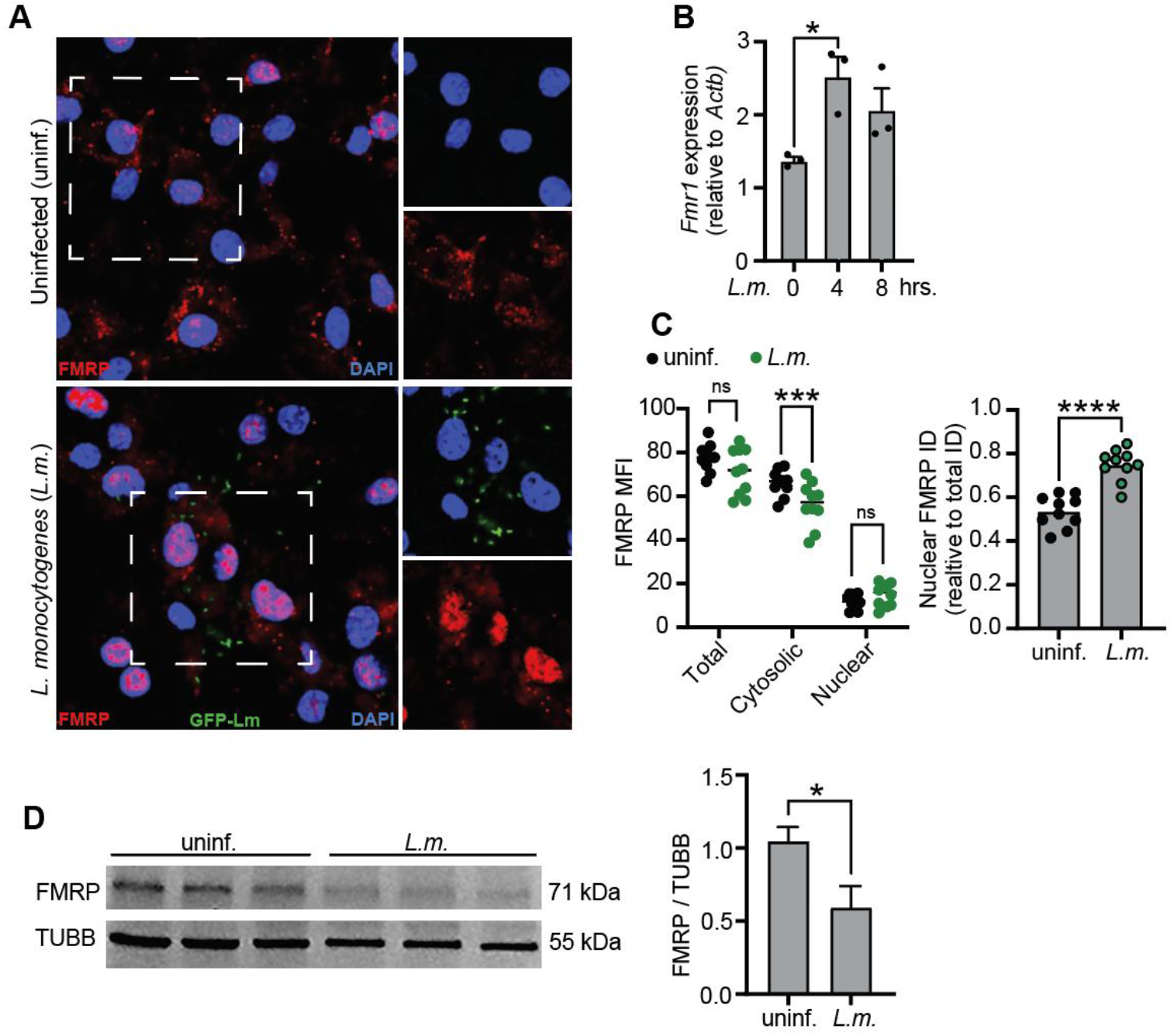
FMRP in murine bone marrow derived macrophages (BMDMs). (A) Immunocytochemistry (ICC) of WT mouse BMDMs infected or not with GFP tagged *L. monocytogenes* at MOI 5 for 4 hours and immunolabeled for FMRP (red). (B) *Fmr1* gene expression in mouse BMDMs infected or not with *L. monocytogenes* for 4 or 8 hours. RT-qPCRs are the mean of 3 replicates ± SD, n = 3 and are representative of at least 2 independent experiments. (C) Mean fluorescence intensity (MFI) and integrated density (ID) quantification of ICC. Each dot represents one random field of view. (D) Representative FMRP immunoblot from macrophages infected or not with *L. monocytogenes* for 4 hours. Immunoblot and ICC images are representative of 3 independent experiments. Statistical significance was determined using a linear regression model (1C, MFI) or exact *p*-value (1B-D) where *, *p* ≤0.05; ***, *p* < 0.005; ****, *p* < 0.0001; ns, not significant.

### *Listeria monocytogenes* infection induces *Fmr1* gene expression

We next wondered if *Fmr1* expression might be similarly upregulated by infection. We again infected mouse BMDMs with *L. monocytogenes* and observed an increase in *Fmr1* gene expression at 4 hours post infection (Fig 1B). In tandem, we also compared FMRP protein levels between infected and uninfected macrophages. No significant difference in total cellular FMRP was observed by immunocytochemistry 4 hours post infection (Fig 1C). However, when we compared cellular compartments, cytosolic FMRP decreased and the ratio of nuclear to total FMRP increased following infection (Figs 1C, S1A). Interestingly, we observed decreased FMRP protein in *L. monocytogenes*-infected macrophages compared to uninfected controls at 4 hours post infection by immunoblot analysis (Figs 1D, S1B). These data indicate that while infection induces *Fmr1* gene expression, FMRP protein is reduced and translocates from the cytoplasm, accumulating in the nucleus upon infection.

### FMRP modulates bacterial induced IL-6

FMRP has been implicated in immune dysregulation whereby loss of FMRP is associated with increased risk of infectious disease (10), however a direct role for FMRP in the regulation of innate immune cytokine production remains unexplored. To investigate the contribution of FMRP to antibacterial innate immune responses, we infected BMDMs derived from *Fmr1*^*+/y*^ (WT) and *Fmr1*^*-/y*^ (KO) mice with *L. monocytogenes*. We measured innate immune transcripts in *L. monocytogenes* infected BMDMs and observed a robust induction of proinflammatory and type I IFN genes (Figs 2A, S2A). While *Ifnb* and *Ifit1* expression was similar between WT and KO macrophages (Fig S2A), *Il6, Tnfa, and Il1b* induction was impaired in macrophages lacking FMRP (Fig 2A). In tandem with *Il6* gene expression, we also measured IL-6 protein levels in *L. monocytogenes*-infected BMDMs. Consistent with transcriptional data, we observed diminished secreted IL-6 in supernatants from FMRP deficient BMDMs (Fig 2B). Together, these results suggest that FMRP contributes to antibacterial proinflammatory gene expression and cytokine production.

**Figure 2:**
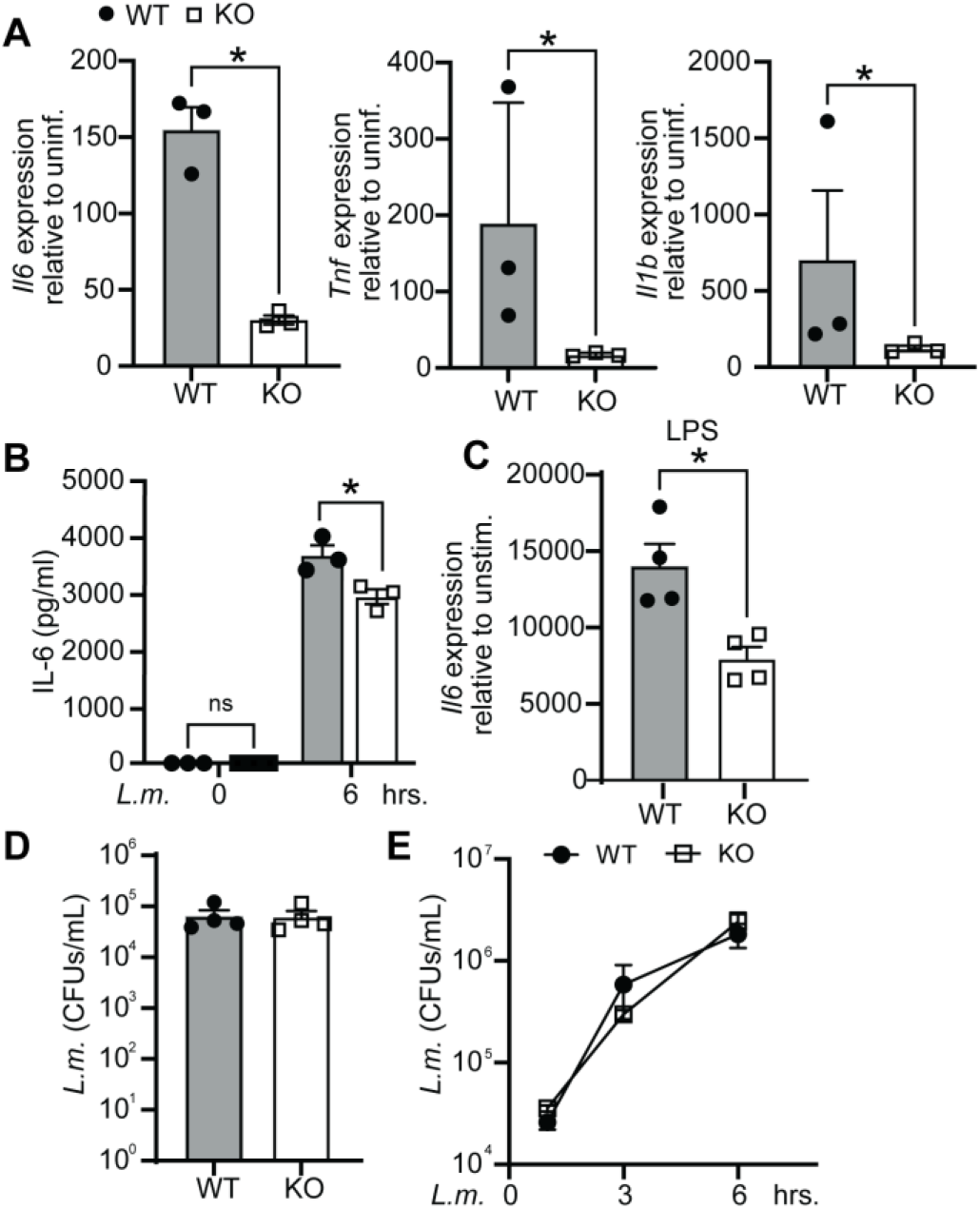
FMRP modulates proinflammatory cytokine production. (A) RT-qPCR in WT and *Fmr1* KO BMDMs infected with *L. monocytogenes* at MOI 10 for 4 hours, assessing *Il6, Tnfa*, and *Il1b*. (B) ELISA of IL-6 in BMDM supernatant after 6 hours of *L. monocytogenes* infection. (C) RT-qPCR of *Il6* in BMDMs following treatment with LPS for 4 hours. (D) Bacterial burden in BMDMs 1 hour after infection with *L. monocytogenes* at MOI of 1. (E) Bacterial burden in WT or *Fmr1* KO BMDMs infected with *L. monocytogenes* over time. RT-qPCRs are the mean of 3 replicates ± SD, n = 3 and are representative of at least 2 independent experiments. Statistical significance was determined using an exact *p*-value where *, *p* ≤0.05; ns, not significant.

### Bacterial uptake and survival are independent of macrophage FMRP

One potential explanation for the defect in IL-6 induction in the absence of FMRP could be reduced susceptibility to bacterial infection. We stimulated BMDMs with 100 ng/ml of LPS for 4 hours to target TLR4 sensing and observed a robust upregulation of *Il6* mRNA transcript levels in both WT and KO macrophages. As in *L. monocytogenes*-infected macrophages, proinflammatory transcript induction was reduced in LPS-stimulated KO cells compared with WT following LPS stimulation (Figs 2C, S2B). This finding suggests that the absence of FMRP perturbs innate immune pathways normally triggered by infection.

To further ensure differences in bacterial burden were not driving the reduced expression of *Il6*, we infected BMDMs with *L. monocytogenes* at an MOI of 1 and measured bacterial uptake at 1 hour post infection and observed no differences in intracellular *L. monocytogenes* in WT or *Fmr1* KO macrophages (Figs 2D, S2C). We again infected WT and *Fmr1* KO BMDMs with *L. monocytogenes* at a MOI of 1, washed cells with gentamycin, and measured intracellular bacterial burden over time. Gentamycin was used to kill extracellular bacterial but does not impact intracellular bacteria. We observed no difference in *L. monocytogenes* replication at 3- or 6-hours post infection (Fig 2E). Thus, we concluded that macrophage FMRP does not impact infectivity or replication of these bacteria.

### Macrophage inflammasome priming is not dependent on FMRP

Altered levels of pro-inflammatory gene expression in infected *Fmr1* KO macrophages may have consequences on cell death pathways, specifically the inflammasome. Indeed, it is known that activation of the canonical inflammasome pathway requires 2 signals: priming followed by activation. To determine if FMRP deficient macrophage viability was impacted by just signal 1, we treated BMDMs with 10 ng/ml of Pam3CSK4, a synthetic mimetic of bacterial lipopeptide found in gram positive and negative bacteria and recognized by TLR2/TLR1. We assessed macrophage membrane permeabilization by staining cells with propidium iodide (PI), a membrane-impermeable dye which fluoresces upon binding to nucleic acids in dead cells. In tandem, we assessed plasma membrane damage as a readout for cytotoxicity by measuring the extracellular release of the cytosolic enzyme lactate dehydrogenase (LDH) into the cell culture media. Pam3CSK4 treatment resulted in a modest induction of cell death which was independent of genotype (Fig 3A-B). We next treated cells with 100 ng/ml of the bacterial endotoxin LPS, a component of gram-negative bacteria recognized by TLR4. Cell survival was also equivalent between LPS-treated WT and KO BMDMs when normalized to baseline levels (Fig 3C), and LPS did not induce LDH release when compared to unstimulated cells (Fig 3D).

**Figure 3.**
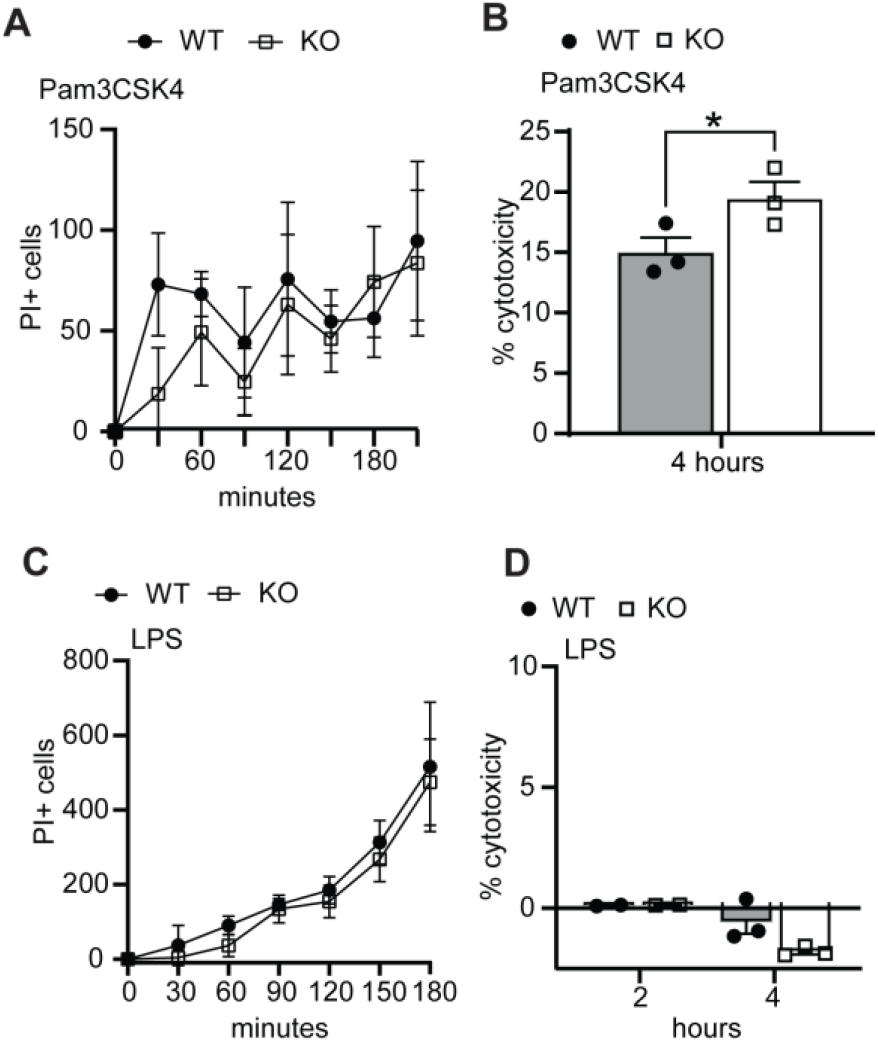
Cell death in primed FMRP deficient bone marrow derived macrophages (BMDMs). (A) Propidium iodide (PI) positive cells in WT and *Fmr1* KO BMDMs primed with 10 ng/ml Pam3CSK4 for the indicated times and normalized to baseline. (B) % cytotoxicity calculated from secreted lactate dehydrogenase (LDH) in WT and *Fmr1* KO BMDMs following Pam3CSK4 stimulation for 4 hours. (C) As in A but with 100 ng/ml LPS. (D) As in B but with LPS stimulation. PI+ cells and LDH levels (cytotoxicity) are the mean of 3 replicates ± SD and are representative of at least 3 independent experiments. Statistical significance was determined using an exact *p*-value where *, *p* ≤0.05.

### FMRP deficiency exacerbates inflammatory cell death

We next interogated whether FMRP contributes to cell survival following both inflammasome signals 1 and 2. To determine if FMRP deficiency impacted the ability of macrophages to activate the inflammasome and induce pyroptosis, we primed BMDMs with 20 ng/ml LPS followed by stimulation with the microbial toxin nigericin to stimulate the canonical inflammasome pathway. FMRP deficient macrophages exposed to nigericin exhibited enhaced PI staining in comparison with WT macrophages at 150 and 210 minutes post infection (Fig 4A, B). In line with this, cytotoxicity was increased in *Fmr1* KO macrophages primed with either LPS or 10 ng/ml Pam3CSK4 followed by nigericin (Fig 4C).

**Figure 4.**
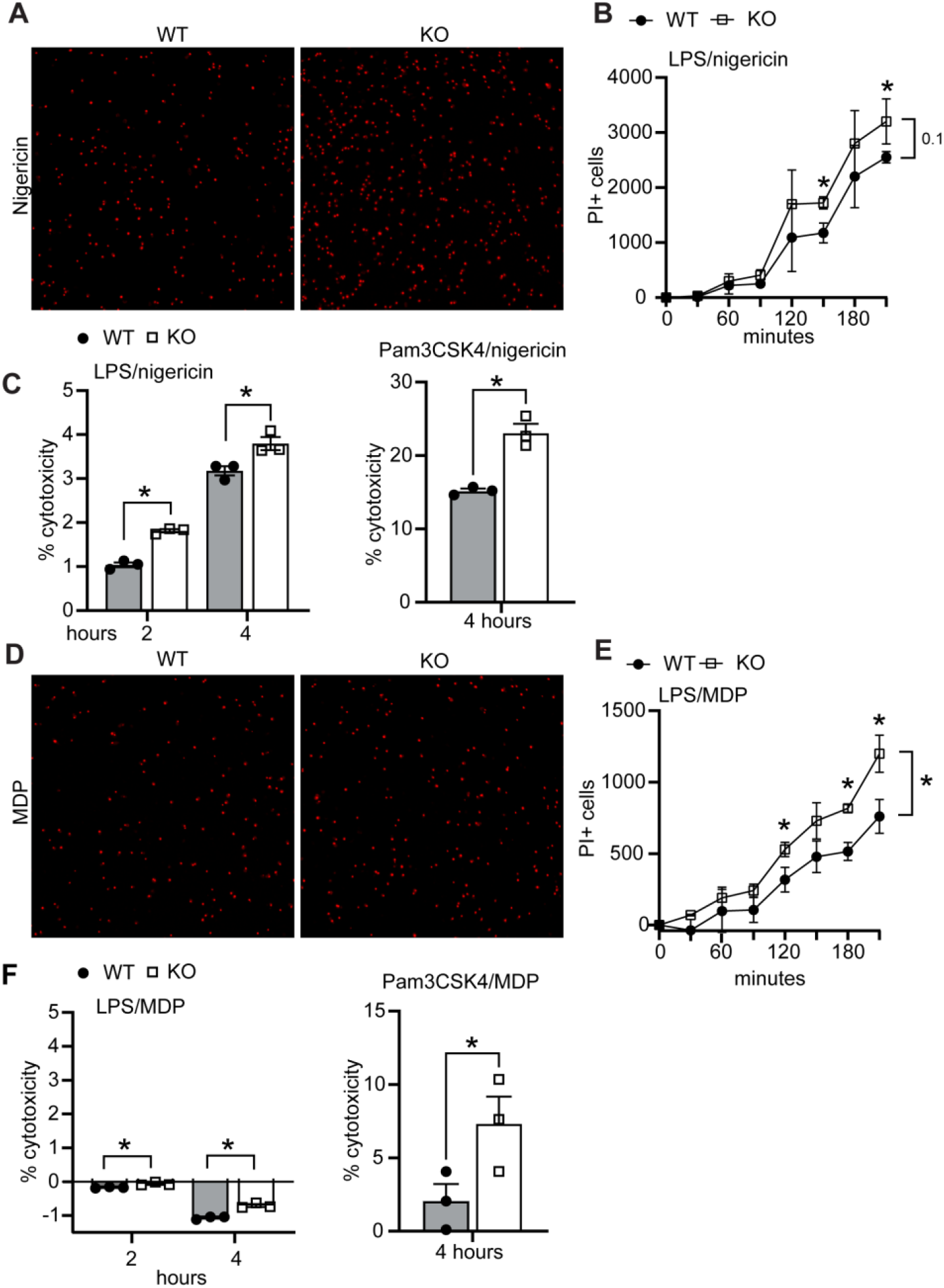
Inflammatory cell death in FMRP deficient bone marrow derived macrophages (BMDMs). (A) Propidium iodide (PI) staining in WT and *Fmr1* KO BMDMs primed with 20 ng LPS for 3 hours followed by 10µM nigericin for 180 minutes. (B) PI+ cells as in A over a 210-minute time course. Each time point is normalized to baseline. (C) % cytotoxicity calculated from secreted lactate dehydrogenase (LDH) in WT and *Fmr1* KO BMDMs primed with LPS or Pam3CSK4 for 3 hours followed by NLRP3 activation with 10µM nigericin. (D) As in A but with NOD2/NLRP3 activation with 25ng/ml MDP for 180 minutes. (E) As in B but with MDP. (F) As in C but with MDP. PI+ cells and LDH levels are the mean of 3 replicates ± SD and are representative of at least 3 independent experiments. Statistical significance was determined using an exact *p*-value where * *p* ≤0.05.

To test whether the increased susceptibility to cytotoxicity was restricted to NLRP3 inflammasome sensing, we switched agonists and stimulated cells with muramyl dippeptide (MDP) to stimulate NOD2/NLRP3 sensing or Poly(dA:dT) to stimulate the AIM2 inflammasome after priming with LPS. MDP stimulation resulted in an increase in cell death in the absence of FMRP (Fig 4D-E). However, LPS priming with MDP stimulation failed to induce LDH release (Fig 4F). Pam3CSK4 priming with MDP stimulation resulted enhanced cytotoxicity in *Fmr1* macrophages (Fig 4F). Poly(dA:dT) transfection resulted in cell death independent of genotype (Fig S3). Together, these data indicate that activated macrophages lacking FMRP are more susceptible to inflammasome activation via different intracellular agonists.

## DISCUSSION

FMRP is highly expressed in neurons and glial cells where it functions as a regulator of proteins critical for proper brain development, synaptic function, and cognition. In addition to its established role in regulating mRNA translation and stability in brain cells, we demonstrate a role for FMRP in macrophages, key cells in innate immunity which act as part of the first line of defense against invading pathogens.

Here, we show that *L. monocytogenes* infection of WT macrophages resulted in upregulation of *Fmr1* gene expression, but that protein levels were reduced. This inverse relationship mirrors that of premutation carriers in which *FMR1* mRNA levels are elevated, in some cases reaching toxic levels resulting in mitochondrial dysfunction, while FMRP protein levels are reduced as a result of impaired translation in premutation carriers (8). The precise cause for the reduction in FMRP following macrophage infection remains to be determined, but may be due to consumption or degradation.

We also show that FMRP primarily localized to the cytoplasm in resting macrophages but relocated to the nucleus upon infection. While FMRP is generally positioned in the cytoplasm in various cell types, it does localize to the nucleus during early embryogenesis and in response to certain cell signals and stimuli. The nuclear functions of FMRP are not well understood, but evidence suggests nuclear FMRP interacts with small nucleolar RNAs to modulate export and translation of target mRNAs (27–29). Given the extent of macrophage membrane and cytosolic receptors activated by *L. monocytogenes*, one possible explanation for FMRP nuclear localization in the context of infection is to modulate immune mRNA export and stability, although this has not been tested. We also demonstrate that the absence of FMRP confers increased susceptibility to inflammatory cell death in macrophages induced by a variety of intracellular receptors. Others have shown protection conferred by overexpressing *Fmr1* in LPS-treated cardiomyocytes or enhanced cell death in FMRP deficient human colorectal cancer tissues (16, 19). However, another study found that the absence of FMRP reduced programmed cell death in neurons in neonatal mice (30), suggesting FMRP invokes tissue-and/or stimulant specific responses.

Here, our data indicate that the loss of FMRP in macrophages resulted in an impaired proinflammatory response to multiple stimuli including bacterial infection. We find that the proinflammatory cytokine IL-6 is reduced in the absence of FMRP in *L. monocytogenes*-infected macrophages. Consistent with (11), we observed no difference in bacterial burden between genotypes in *L. monocytogenes*-infected BMDMs, pointing to a more direct role in FMRP modulation of proinflammatory cytokine production. While studies have reported elevations in neonatal mouse brain cortical tissue or astrocyte IL-6 protein levels in *Fmr1* KO models (31), we observed no baseline differences in proinflammatory and antiviral cytokine genes in primary macrophages, consistent with (18, 32). FMRP has been suggested to regulate IL-6 and TNF-α levels (33). Though several studies have reported elevated proinflammatory gene expression and protein levels following LPS treatment (18, 34), we observed a reduction in *Il6* mRNA and protein levels in *Fmr1* KO macrophages, both those infected with *L. monocytogenes* or stimulated with LPS (Fig 2). One potential explanation for this discrepancy is differences in background strain and experimental approach; both (18) and (31) used FVB.129P2-Pde6b+ Tyrc-ch Fmr1tm1Cgr/J mice and (18) used an *in vivo* approach examining hippocampal regions. In contrast, this study used cells cultured from B6.129P2-Fmr1tm1Cgr/J mice. Indeed, several studies have observed strain dependent differences in the FXS mouse model (35, 36). This incongruity may also point to cell specific differences as elevations in IL-6 secreted protein levels were similarly reported in primary FMR1 KO astrocytes treated with LPS (34), while we used BMDMs. The potential for cell specific genotypic differences in inflammatory responses is further supported by another study in microglia which found equivalent levels of IL-6 and TNF-α levels following treatment with LPS (32).

Taken together, these data expand existing evidence indicating innate immune dysregulation in the absence of FMRP and show an impaired proinflammatory response to bacterial infection, which has public health implications for individuals with FXS. Collectively, the results shown here support a role for FMRP in modulating proinflammatory gene expression and inflammatory cell death in macrophages.

## Acknowledgements

This work was supported by resources provided by the TNBRC Base Grant P51OD011104 (RRID:SCR_008167) and TNBRC Confocal Microscopy and Molecular Pathology Core (RRID: SCR_024613).

## Author Contributions

Conceptualization, KJV; data curation and formal analysis, BNM, LAH and KJV; funding acquisition, KJV; investigation, BNM and KJV; methodology, CGW, ROW, KJV; project administration KJV; resources, KJV; supervision, ROW, KJV; validation, BNM, TF, KJV; visualization, BNM, CGW, KJV; writing – original draft, BNM, KVJ; writing – review and editing, BNM, CGW, TF, ROW, LAH KJV.

