## Supplemental Figures 1-3 for "Fragile X messenger ribonucleoprotein modulates IL-6 induction and inflammatory cell death in macrophages"

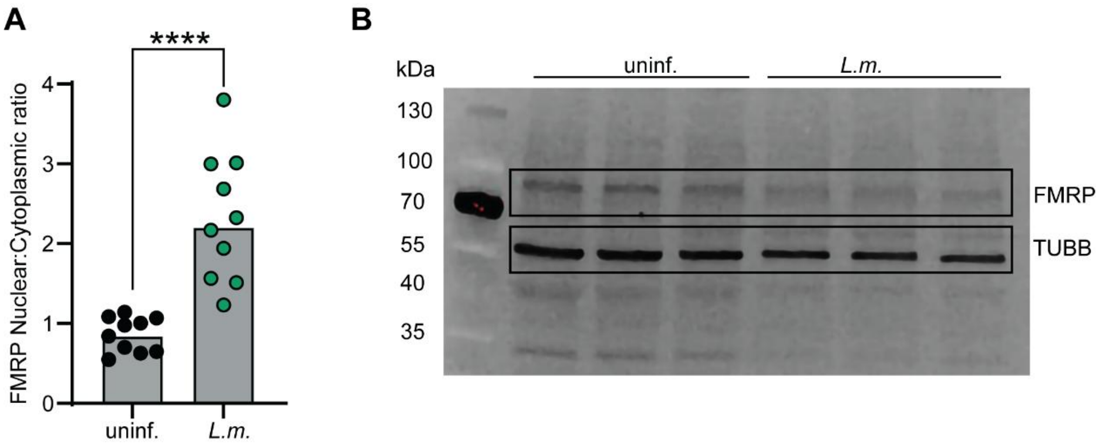


**Figure S1: FMRP in murine bone marrow derived macrophages (BMDMs).** (A) Nuclear: cytoplasmic ratio of FMRP integrated density quantified by immunocytochemistry in murine BMDMs uninfected or infected with *L. monocytogenes* for 4 hours. Each dot represents a random field of view. Statistical significance was determined using an exact *p*-value where ****, $p < 0.0001$. (B) Larger area of FMRP immunoblot shown as a cropped image in Figure 1D from macrophages infected or not with *L. monocytogenes* for 4 hours.

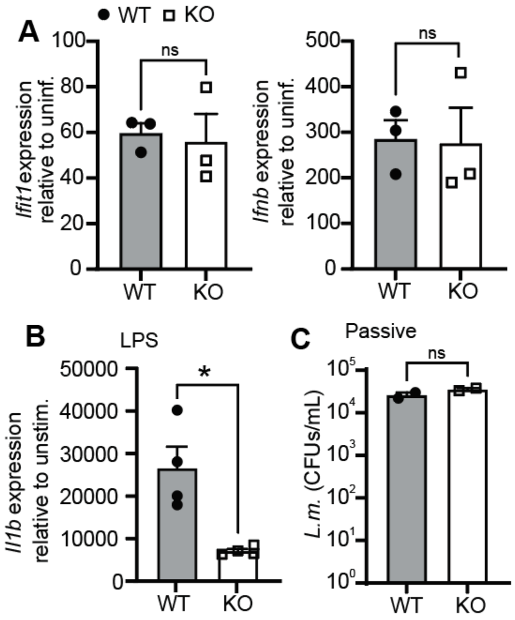


**Figure S2: FMRP modulates proinflammatory cytokine production.** (A) RT-qPCR in WT and *Fmr1* KO BMDMs infected with *L. monocytogenes* at MOI 10 for 4 hours, assessing *Ifit1* and *Ifinb*. (B) RT-qPCR in WT and *Fmr1* KO BMDMs stimulated with 100 ng/ml LPS for 4 hours, assessing *Il1b*. (C) Bacterial burden in BMDMs 1 hour after passive (uncentrifuged) infection with *L. monocytogenes* at MOI of 1. RT-qPCRs are the mean of 3 replicates ± SD, n = 3 and are representative of at least 2 independent experiments. Statistical significance was determined using an exact *p*-value where *, $p \leq 0.05$; ns, not significant.

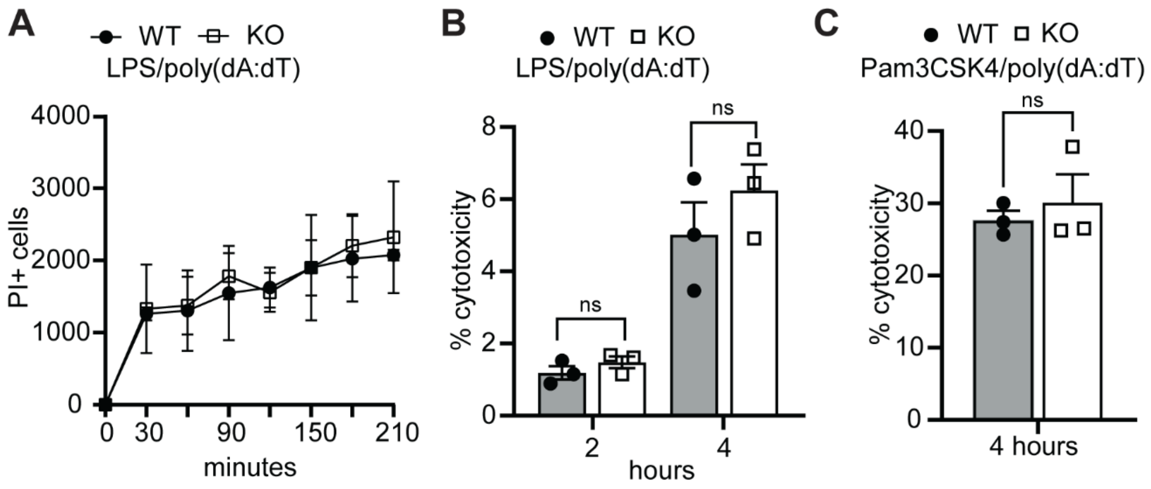


**Figure S3. Inflammatory cell death in FMRP deficient bone marrow derived macrophages (BMDMs).** (A) Propidium iodide (PI) positive WT and *Fmr1* KO BMDMs primed with 20 ng/ml LPS for 3 hours followed by AIM2 activation with 1μg/ml Poly(dA:dT) for the indicated times. (B) % cytotoxicity (secreted LDH) in BMDMs primed with 20 ng/ml LPS for 3 hours followed by AIM2 activation with poly(dA:dT). (C) As in B but primed with 10 ng/ml Pam3CSK4. Experiments are the mean of 3 replicates ± SD and are representative of at least 3 independent experiments. Statistical significance was determined using an exact *p*-value; ns, not significant.
